# Dynamic causal modeling of layered magnetoencephalographic event-related responses

**DOI:** 10.1101/2020.07.20.208504

**Authors:** Stephan J. Ihle, James J. Bonaiuto, Sven Bestmann, Klaas Enno Stephan, Gareth R. Barnes, Jakob Heinzle

**Affiliations:** Laboratory of Biosensors and Bioelectronics, Institute for Biomedical Engineering, University of Zurich & ETH Zurich, Zurich, Switzerland; Translational Neuromodeling Unit (TNU), University of Zurich & ETH Zurich, Zurich, Switzerland; Institut des Sciences Cognitives Marc Jeannerod, CNRS UMR 5229, Bron, France; Université Claude Bernard Lyon 1, Université de Lyon, France; Department for Clinical and Movement Neuroscience, UCL Queen Square Institute of Neurology, University College London, London, UK; Wellcome Centre for Human Neuroimaging, UCL Queen Square Institute of Neurology, University College London, WC1N 3BG London, UK

**Keywords:** Cortical Layers, MEG, Dynamic Causal Modeling, Canonical Microcircuit, High-resolution MEG

## Abstract

The layered architecture of cortex is thought to play a fundamental role in shaping cortical computations. However, direct electrophysiological measurements of layered activity are not possible non-invasively in humans. Recent advances have shown that a distinction of two layers can be achieved using magnetoencephalography in combination with head casts and advanced spatial modeling. In this technical note, we present a dynamic causal model of a single cortical microcircuit that models event related potentials. The model captures the average dynamics of a detailed two layered circuit. It combines a temporal model of neural dynamics with a spatial model of a layer specific lead field to facilitate layer separation. In simulations we show that the spatial arrangement of the two layers can be successfully recovered using Bayesian inference. The layered model can also be distinguished from a single dipole model. We conclude that precision magnetoencephalography in combination with detailed dynamical system modeling can be used to study non-invasively the fast dynamics of layered computations.

## Introduction

Its layered structure is one of the defining anatomical features of cerebral cortex (Douglas and Martin, 2004). How this layered, hierarchical organization relates to the computational properties of the human brain is still not understood in detail. However, theoretical predictions have been made about a possible role of the layered architecture (Bastos et al., 2012; Douglas and Martin, 2007; Heinzle et al., 2007). For example, predictive coding (Friston, 2005; Rao and Ballard, 1999), a specific instantiation of the general “Bayesian brain” theory (Knill and Pouget, 2004), can be implemented by a hierarchy of layered microcircuits, with distinct computational roles of neurons in different layers (Bastos et al., 2012; Shipp, 2016; Stephan et al., 2019).

In order to test hypotheses like predictive coding in humans, non-invasive measurements of layered activity are required. One option for this is presented by recent advances in high-resolution layered functional magnetic resonance imaging (fMRI) (De Martino et al., 2015; Kok et al., 2016; Muckli et al., 2015). However, fMRI provides an indirect and intrinsically slow measure of neural activity which might suffer from blood draining effects when applied to layers (Heinzle et al., 2016). Separation of feedforward (bottom-up) and feedback (top-down) streams would be facilitated by a more direct, electrophysiological measurement of layered activity as, for example, provided by magnetoencephalography (MEG, for a review see Baillet, 2017). The spatial specificity of MEG can be improved by assuring precise positioning of the head within the MEG and a high-resolution cortical model of layers (Bonaiuto et al., 2018a; Troebinger et al., 2014b). Simulation studies have shown that under these conditions oscillatory activity can be reliably distinguished between two layers located at the pial surface or gray matter - white matter boundary, respectively (Bonaiuto et al., 2018b; Troebinger et al., 2014a). To date, these studies have focused on inversion of a precise “spatial model” describing the mapping from one (or several) current dipole(s) distributed on the two cortical surfaces to sensor activity.

In contrast, temporal models which describe the dynamics of neural activity are routinely considered within the dynamic causal modeling (DCM) framework. DCMs of EEG and MEG are generative models that explain measured MEG/EEG sensor data through a forward mapping of activity in one or several connected cortical columns. These are modeled as microcircuits consisting of several neural populations (e.g. supra- and infragranular pyramidal neurons, spiny stellate and inhibitory neurons of a cortical column). For reviews see for example (Kiebel et al., 2009; Moran et al., 2013). However, the spatial model in DCM has, to date, not allowed the investigation of neural signals at the resolution of layers. Instead, this approach has considered contributions of individual populations from different layers by attributing the weighted summed activity to a single point source or current dipole. Using this forward model, networks of layered microcircuits were used to study the relative importance of feed-forward and feed-back connections in auditory mismatch (Garrido et al., 2008) and the processing of face (Chen et al., 2009) and somatosensory stimuli (Auksztulewicz and Blankenburg, 2013). Recently, a dynamic causal model was applied to invasively recorded depth dependent cortical activity of mice (Pinotsis et al., 2017), directly predicting layered signals. This model was based on previous work that proposed a small network of groups of neurons to explain somatosensory evoked potentials (Jones et al., 2007) and mu-rhythms (Jones et al., 2009) in MEG. Notably, Jones et al (2007) used detailed compartmental neurons to directly model currents in dendritic trees. They manually adjusted parameters to match the resulting current dipole changes to source reconstructed MEG data.

Here, we augment previous generative modeling attempts that focus mainly on temporal models (e.g. Garrido et al., 2008) or on spatial aspects (Bonaiuto et al., 2018a; Troebinger et al., 2014a). We fit a temporal model of a laminar circuit to simulated MEG data while making use of precise anatomical information. This allows us to assign different dipoles to different cortical layers. As a basis for the temporal model we use a canonical microcircuit (Bastos et al., 2012; Moran et al., 2013) approximation to the Jones model (Jones et al., 2007). In simulations, we show that a model with correct layer information can be distinguished from models where this information is reversed or missing at signal to noise levels that are commonly observed in MEG. We show how the addition of temporal information removes considerable ambiguity in the spatial model by constraining sources to lie on the correct sulcal wall. We then investigate how this inference on layered circuits depends on parameter settings and how well the distance between current dipoles in supra- and infragranular layers can be estimated. Finally, we investigate to what degree adding more sensors increases the sensitivity of the method. In summary, our simulations show that inversion of layered dynamic models is possible under reasonable SNR settings.

## Methods

The simulations and model inversions performed in this paper use the DCM for EEG and MEG framework (David et al., 2006; Kiebel et al., 2009, 2006). The generative model for MEG data consists of a temporal model (a set of differential equations that describe the temporal evolution of neural activity in a layered microcircuit) and a spatial model (the lead field mapping that converts current dipoles in different cortical layers to signals in MEG sensors).

We first introduce the dynamical layered model, a canonical microcircuit (Bastos et al., 2012; Moran et al., 2013) which we adapted to replicate the dynamics of a detailed compartmental neuronal circuit used to explain somatosensory event related MEG responses (Jones et al., 2007), referred to as Jones compartmental microcircuit model (JCM) throughout this paper. Second, we explain the spatial model used for simulating MEG traces from layered cortical activity (Bonaiuto et al., 2018b, 2018a; Troebinger et al., 2014a, 2014b). Third, we outline the simulation and inversion of DCM for layered MEG and present the simulations carried out to test the feasibility of inferring layered circuit activity from MEG data. Figure 1 graphically summarizes the model used in this paper.

**Figure 1:**
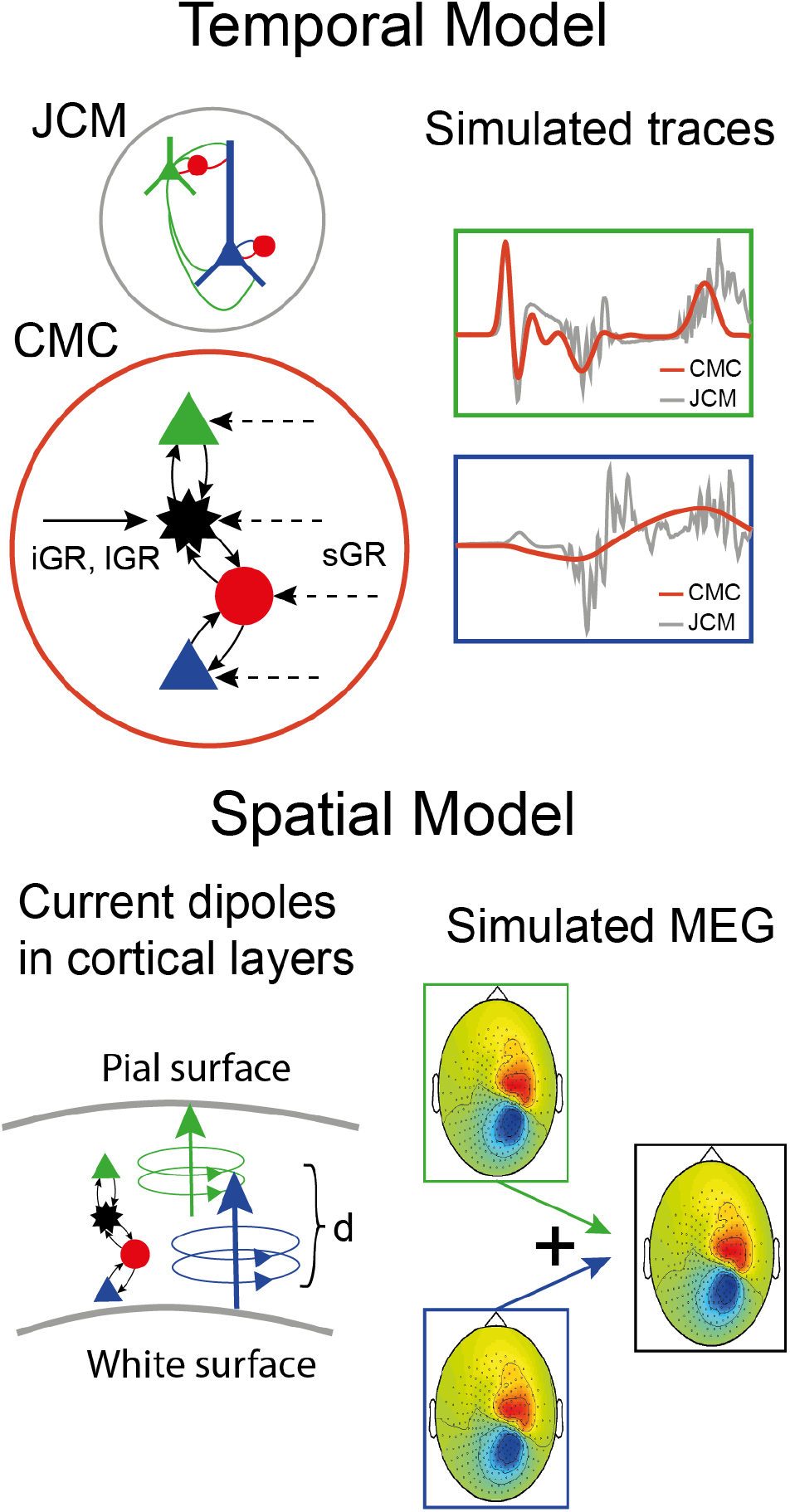
Illustration of the generative model for layered MEG. *Top:* The temporal dynamics of the model were controlled by a canonical microcircuit model (CMC: spiny stellates: black, inhibitory interneurons: red, superficial pyramidal cells: green, deep pyramidal cells: blue). The CMC received two waves of bottom-up input to the granular layers (iGR and lGR) and top-down input to the pyramidal cells (sGR). The parameters of the CMC were fitted in order to match responses of CMC supra- and infra-granular pyramids (red traces) to the corresponding cells simulated by the JCM red traces. *Bottom:* In order to generate MEG sensor signals we assumed that the activity of the pyramidal neurons generated a current dipole positioned in the upper (pial surface) and lower (grey matter – white matter boundary) layer of cortex, with an orientation perpendicular to the cortical surface. The distance *d* specifies the distance between the two dipoles and hence relates to cortical thickness.

### The temporal model

In this section, we describe the canonical microcircuit model (MCM; Moran et al., 2013) and how it was initially fitted to the Jones compartmental microcircuit model (Jones et al., 2007).

### Canonical microcircuit model

The layered microcircuit in this paper was a neural mass model (Moran et al., 2013) that describes the dynamics of a cortical layer by focusing on key neuronal populations in different laminae. In order to allow for compatibility with the standard models in SPM, we used the canonical microcircuit model (CMC) as the basic layered neuronal network. This circuit includes one population of spiny stellate neurons, one population of inhibitory interneurons, and two populations of pyramidal neurons in layers 2/3 and 5, respectively. A schematic of the model circuit is shown in Figure 1. The dynamics of the populations are controlled by a second order differential equation following the work by Jansen and Rit (1995):

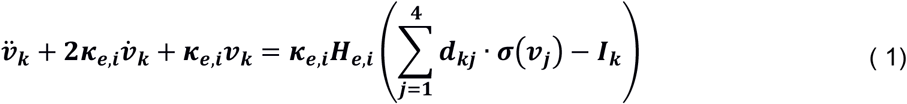

Here, *ν_k_* is the activity of population k. *κ_e,i_* controls the oscillatory behavior of the cell population and may differ between excitatory (e) and inhibitory (i) neurons. *H_e,i_* is a gain parameter defining the maximal postsynaptic potential, *d_kj_* are connectivity parameters and *σ*(*ν_j_*) is a sigmoid function transforming activation into a “synaptic” input. Inhibitory connections are defined by setting the corresponding *d_kj_* to be negative. Finally, *I_k_* is the “external” input into population k.

The parameters of the CMC were adapted to approximate the responses of the JCM. In order to adequately reproduce the JCM with the CMC, we included the three different inputs proposed in (Jones et al., 2007): an initial granular (iGR) layer input targeting layer 4, a supra-granular (SGR) layer input – modelling feedback from higher regions and targeting all four populations, but with different weights (see Table 2, below), and a late granular (lGR) layer input targeting layer 4. All three inputs were described by Gaussian distributions. Table 1 summarizes the corresponding mean values and dispersion.

**Table 1:** Input timing parameters. Inputs were given as Gaussians in time. Mean and standard deviation of the three inputs are given.

| *Input name* | *Input timing (Mean)* | *Input dispersion (STD)* |
| --- | --- | --- |
| iGR | 25 ms | 2.5 ms |
| SGR | 70 ms | 6 ms |
| IGR | 135 ms | 7 ms |

**Table 2:**
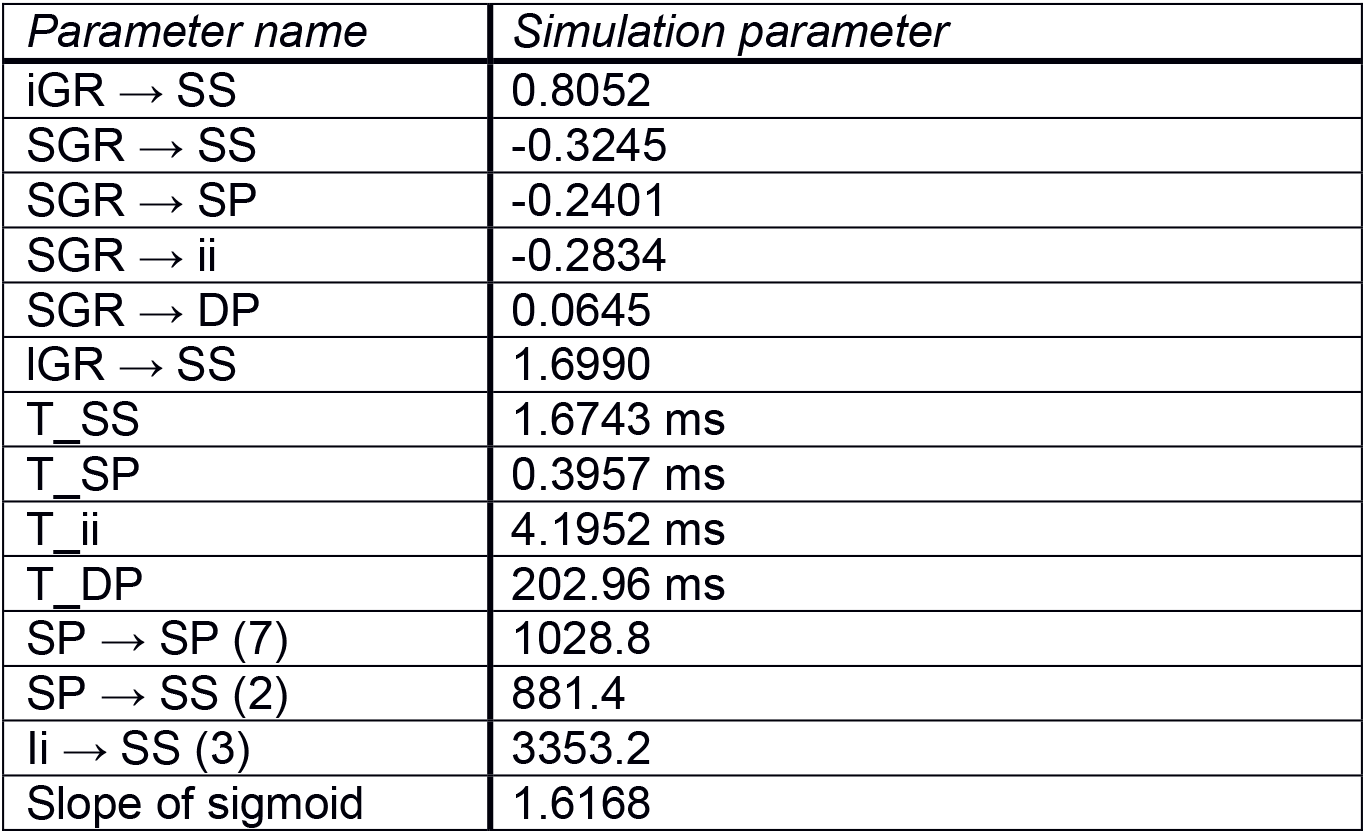
Posterior estimates. The MAP estimates of all fitted parameter of the CMC. The traces were fitted to the JCM. All parameters not listed were kept identical to the SPM implementation of the CMC.

### Fitting the CMC to the JCM

In order to simulate realistic traces of supra- and infragranular currents, we used the model of (Jones et al., 2007). We downloaded the version available at http://senselab.med.yale.edu/senselab/modeldb/ and simulated a somatosensory evoked potential. The model was run 100 times as proposed by Jones et al. (2007). Traces were created by taking the average of these 100 runs. In the next step, we fitted (using SPMs spm_nlsi_GN.m) the CMC to the JCM using a simple forward model that matched the activity of the supra- and infra-granular layer to the corresponding current dipoles of the JCM model. The signal was simulated for the first 160 ms after stimulation (i.e. 0-160 ms). Figure 2A illustrates the traces of the JCM model and the fit. We used these maximum a posteriori (MAP) estimates of this fitting procedure for all subsequent simulations of MEG data. The resulting parameters for the simulation of the temporal model are summarized in Table 2.

**Figure 2:**
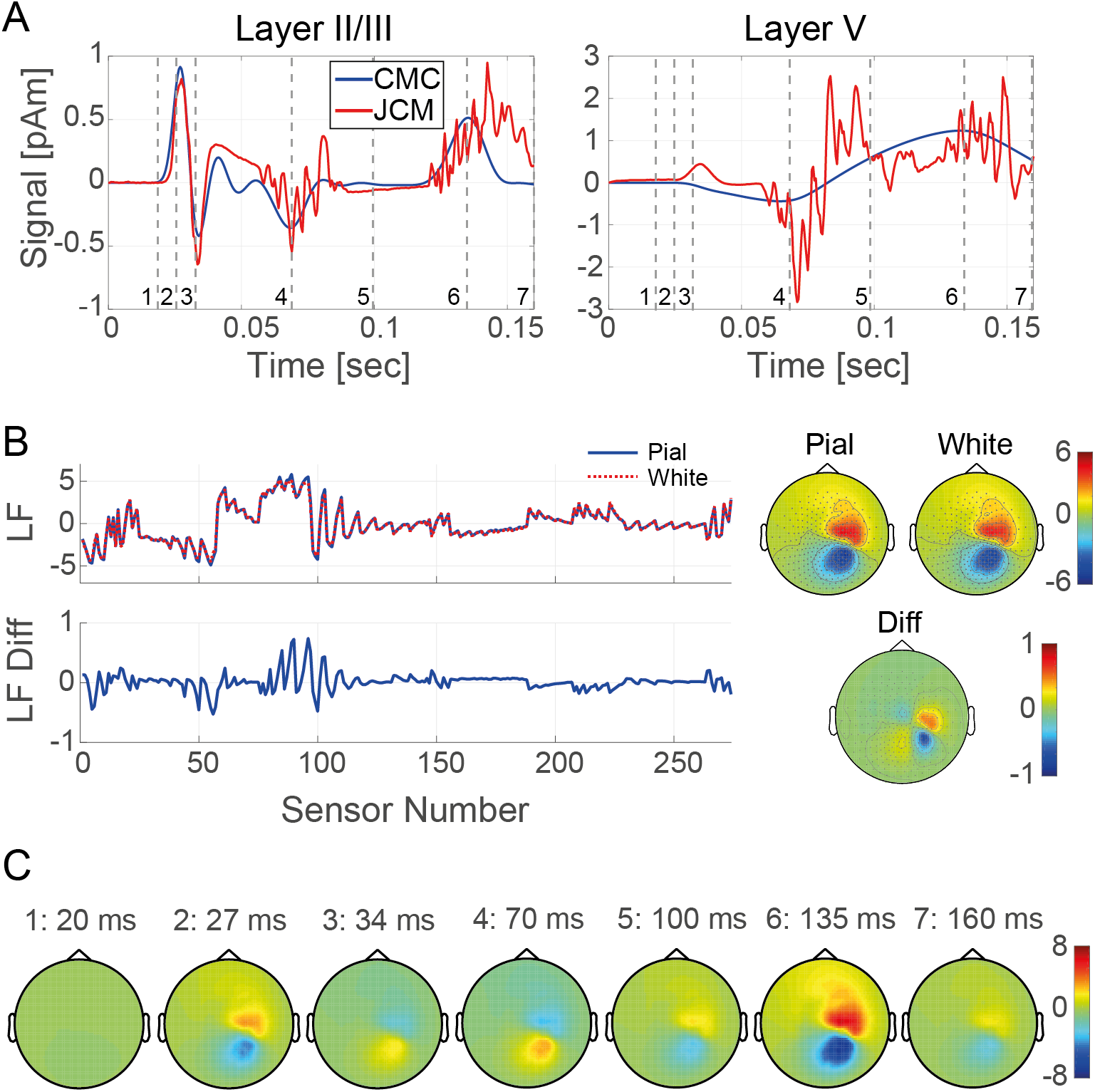
Illustration of MEG sensor activity. **A:** Simulated activity traces of the superficial (Layer II/III) and deep (Layer V) pyramidal neurons based on JCM (red) and corresponding approximated CMC traces (fitted to JCM). Signal strength is given in pAm as in (Jones et al., 2007). Vertical grey dashed lines indicate time points for which brain topographies are illustrated in **C**. **B**: Illustration of lead field. *Left:* Strength of lead field (LF) for two current dipoles on the pial (blue solid) and white matter (red dotted) surfaces at a distance of 4mm at all sensors. The difference between the two curves is plotted below. *Right:* Topography illustrating the same lead fields. **C:** Simulated spatio-temporal traces obtained when the CMC traces in **A** are mapped to the sensors through the lead fields illustrated in **B**. Note that for display purposes the distance between the dipoles was 4 mm in these simulations.

### Spatial Model

While the temporal model fully determines the traces of the two pyramidal population, a spatial model is needed to transform the neuronal signals to sensor data. The spatial model constitutes the forward part of the generative model, mapping hidden states to observations (measurements). The current dipoles of the two populations were modelled as a dipole pair, one for the superficial and one for the deep layer traces. The apical dendrites of both superficial and deep pyramidal neurons are oriented perpendicularly to the cortical surface. Therefore, we assume that the dipole pairs have the same orientation and their positions differ only along this axis (see Bonaiuto et al., 2019, for a comparison of different options to define dipole orientation). In order to mimic a somatosensory stimulation experiment, the dipoles were placed in area BA3b of a normalized brain (MNI space). Here, we used the average location of BA3b according to Papadelis et al (2011). For simulations, dipole orientation was defined by the angle between the dipole and its location vector with respect to the MEG volume conductor origin. Note that in an inversion with real data, the orientation of the dipole would be defined perpendicularly to cortex based on the anatomy of the subject. Here, the radial angle was chosen to be 51° except for simulations where the angle was varied. These exceptions are indicated in the Results section. Finally, the dipoles of the two layers were separated by a distance of 2 millimeters along the dipole orientation. The distance of 2 mm is motivated by cortical thickness measurements in humans which ranges from 1 to 4.5 mm (Fischl and Dale, 2000) but tends to be relatively thin, around 2 mm, in primary sensory areas (Scholtens et al., 2015).

The lead fields of the different layers were calculated with the single shell model (Nolte, 2003) of SPM (Litvak et al., 2011). Synthetic sensor data of an MEG scanner were simulated as described in an earlier paper (Bonaiuto et al., 2018b). Briefly, we used an affine spatial transformation from MNI space to sensor space in order to place the current dipoles in the correct position within the MEG scanner. We then simulated a total of 274 sensors. Figure 2B illustrates the lead fields for two dipole pairs in right BA3b. For display purposes and to highlight the difference between the lead fields of superficial and deep pyramidal neurons, the distance between the two dipoles of a pair was set to 4mm. As expected, the lead fields of the superficial and deep dipole within a pair are very similar. Nevertheless, the two dipoles of a cortical column show differences which are better visible when plotting the lead field in one dimension (Figure 2B). In this work, we try to exploit these small differences to make statements about layer differentiation.

Figure 2C illustrates an example of simulated data from the whole model, using a layered microcircuit in the right BA3b.

### Simulations

In order to test whether the proposed layered circuit could be inferred from sensor data, we simulated MEG data using the generative model described above with a forward model consisting of two lead fields for the two current dipoles of upper and lower layer pyramidal cells. The strategy of the simulations is explained in Figure 3. In general, we simulated data from a model that assigned superficial activity to a current dipole close to the pial surface and activity of deep pyramidal neurons to a current dipole close to the gray-white matter boundary. We refer to this model as the layer correct (LC) model. The sensor traces generated from these simulations were then used as data for subsequent model inversion: In order to investigate whether and how well the assignment of the two sources could be inferred from the data, we inverted several models using the generated data. We then used Bayesian model comparison to establish which was the most likely model, given the data. Model inversion was performed using the variational Bayes approach in SPM (Variational Laplace) which yields the variational negative free energy (F) as an approximation to log model evidence (Friston et al., 2007). Log model evidence is a measure of model goodness used to score different models that were inverted based on the same data set. A difference in log model evidence larger than 3 (equivalent to a Bayes factor larger than 20) is usually considered strong evidence in favour of one model compared to another (Kass and Raftery, 1995).

**Figure 3:**
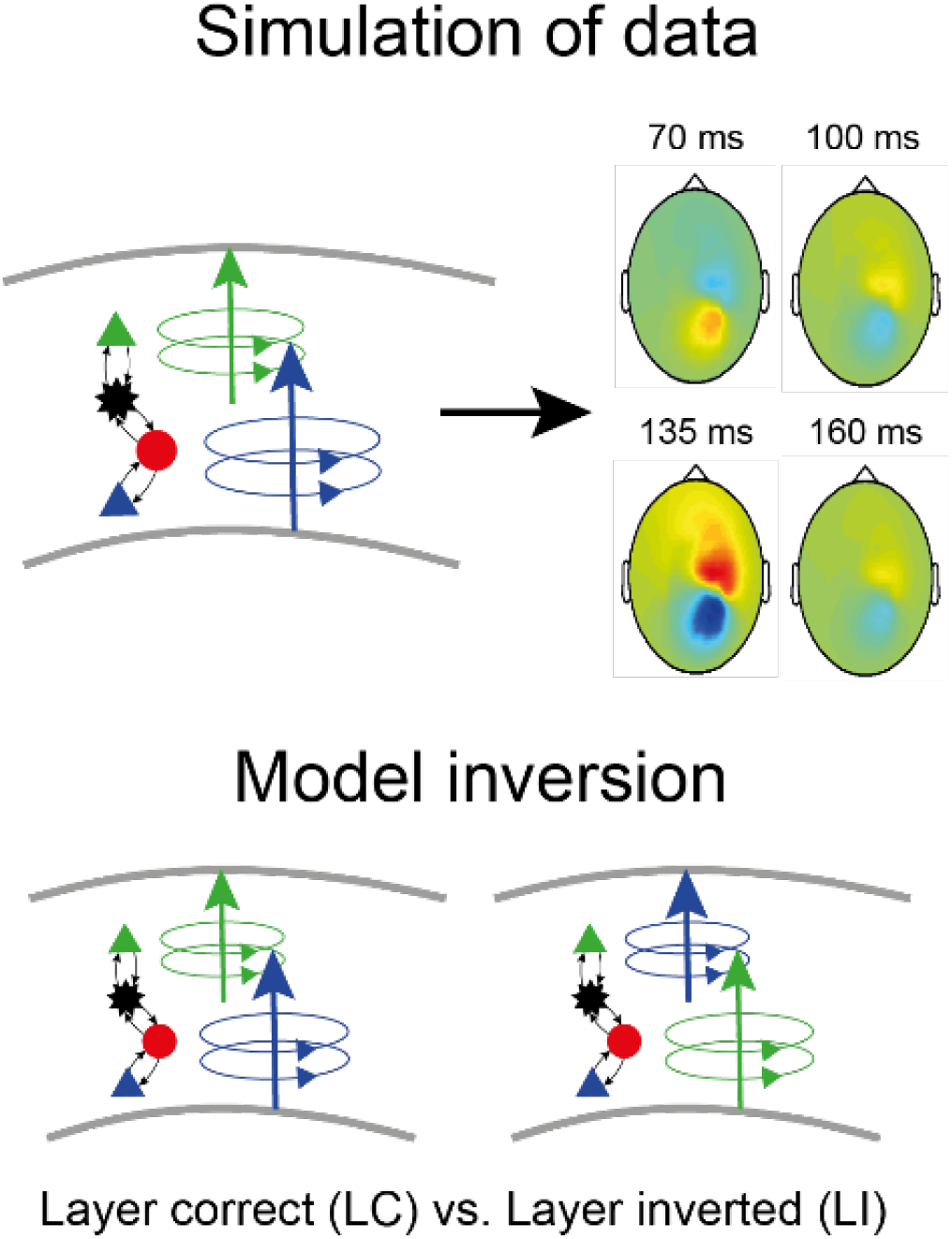
Illustration of simulation strategy. In all simulations, we created models with the correct association between neural populations and dipoles. For model inversion, different models were fitted to the generated data and then compared, for example the layer correct to the layer inverted model.

In a first set of simulations, we compared the LC model to two competing hypotheses: First, we assessed a model with the inverted assignment of layered activity (layer inverted model, LI), that is, with superficial activity assigned to a current dipole close to the gray-white matter boundary and activity of deep pyramidal neurons assigned to a dipole close to the pial surface. Second, we also considered a model where both dipoles were located at the same location (at mid-cortical depth, comparable to traditional DCMs). We refer to this model as layers detached (LD) or single-source model.

After this first set of model comparisons, we conducted a series of simulations to test how sensitive the separation between LC and LI models was and how much it depended on the particular choice of parameters. For this, we simulated different dipole pairs and tested how well the LC and LI models could be separated. In particular, we varied the orientation of the dipoles, cortical thickness, and the distance to the closest sensor. Finally, we conducted a simulation to explore the effects of the arrangement of MEG sensors with a particular focus on the possibility of reducing the number of sensors.

All data were simulated using the CMC with the parameters indicated in Table 2. If not stated otherwise, the model generating the data had a distance of 2 mm between the superficial and deep dipole. After creating the sensor data (sampled at a frequency of 2400 Hz), white Gaussian noise was added to the signals. The noise of each sensor was assumed to be independent from the other sensors and also independent between time points. Furthermore, we assumed the same amount of noise, i.e. variance, for all sensors. We characterize the amount of noise by the signal to noise ratio (SNR) *ξ* as defined by Goldenholz et al (2009):

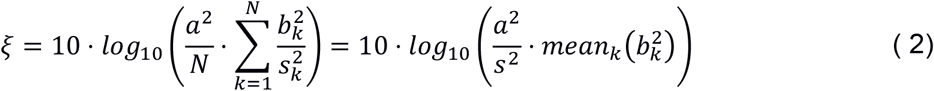

where 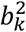 is defined as the signal on sensor k for unit input, 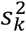 is the variance of the noise added to the k^th^ sensor, N is the number of sensors, and a^2^ the source amplitude. Note that *ξ* describes the overall SNR. The SNR of individual sensors varies and depends on their respective signal strength. Here, we explored SNRs between −20 and 10 dB. In order to avoid effects of any particular noise instance, each simulation was repeated with 20 randomly created noise traces. It is worth noting that the model we propose here can be fitted to averaged event-related responses (ERPs). Thus noise levels have to be compared to the SNR of averaged evoked potentials. Averaging the signal over n trials reduces the variance by a factor of n: 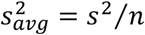. This results in an additive increase of SNR by 10 log_10_ *n*.

### Software note

All code used here was implemented in MATLAB (MathWorks, Inc., Natick, Massachusetts, United States). Simulations were executed either on a personal computer (using Matlab Release 2016b) or on a high-performance computer cluster (EULER, using Matlab Release 2015a) of ETH Zurich, Switzerland. We employed the DCM implementation in SPM12 (release 7219) for general DCM steps and custom written Matlab functions for layered MEG specific parts. The code of all simulations and to generate the figures used in the paper is available on https://gitlab.ethz.ch/tnu/code/dcm-of-layered-meg.

## Results

In this section, we will first show that Bayesian model comparison can distinguish the correctly layered model from two alternative models, one with both dipoles placed in the middle of cortex and one with an inverted dipole position. Next, we explore to what extent model inversion provides model evidence estimates specific enough to distinguish different cortical depths. We then proceed to a thorough analysis of the influence of dipole properties such as orientation and separation of the two layers on model inversion. Finally, we illustrate through simulation how the combined spatio-temporal model is able to distinguish the sign of the current dipole, i.e. the orientation of pyramidal neurons in cortex.

### Proof of Concept

Subsequently, we will show that using both temporal and spatial features improves model prediction at moderate SNR. We do this by comparing the free energy of three different models that were all fitted to the data generated with a layer correct model with a distance of 2 mm between the two layers. The first model incorporated the same dipole locations used for the data generation (layer correct (LC) model). The second model did not use any spatial information about layers (layer detached (LD) model). It located both dipoles at the same position in the middle of cortex, effectively letting them behave as a single dipole. The third model used the same locations as for the LC model but changed the positions of the superficial and deep dipole (layer inverted (LI) model). Thus, this model assigned the time-series of superficial activity to the current dipole in deep layers and vice versa.

By comparing the free energy difference between the LC and the LD model, we can establish to what degree modelling improves when using spatial information about the layer origin as compared to assuming one single spatial source per microcircuit. The comparison of the LC with the LI model shows to what degree different types of layered arrangement can be separated. The difference in spatial information is larger in this second scenario, which results in a clearer distinction based on model evidence. This comparison illustrates how the temporal model can be used to augment spatial information. The results of this model comparison are plotted in Figure 4 for different SNRs ranging from 10dB to −20dB. As expected, models could be more clearly distinguished with increasing SNR. Also, as expected, this difference in model evidence was greater for the more spatially distinct pairings. That is, there was a greater difference between layer correct (LC) and layer inverted (LI) models than between layer correct and layer detached (LD or point-source) models. The improvement was decisive (ΔF > 3) for both comparisons for SNR levels of 0dB or larger. The correct model was reliably selected in most cases even for SNRs of −3.3dB (for some instantiations of noise even −6.7dB) when comparing the LC- and LI-model.

**Figure 4:**
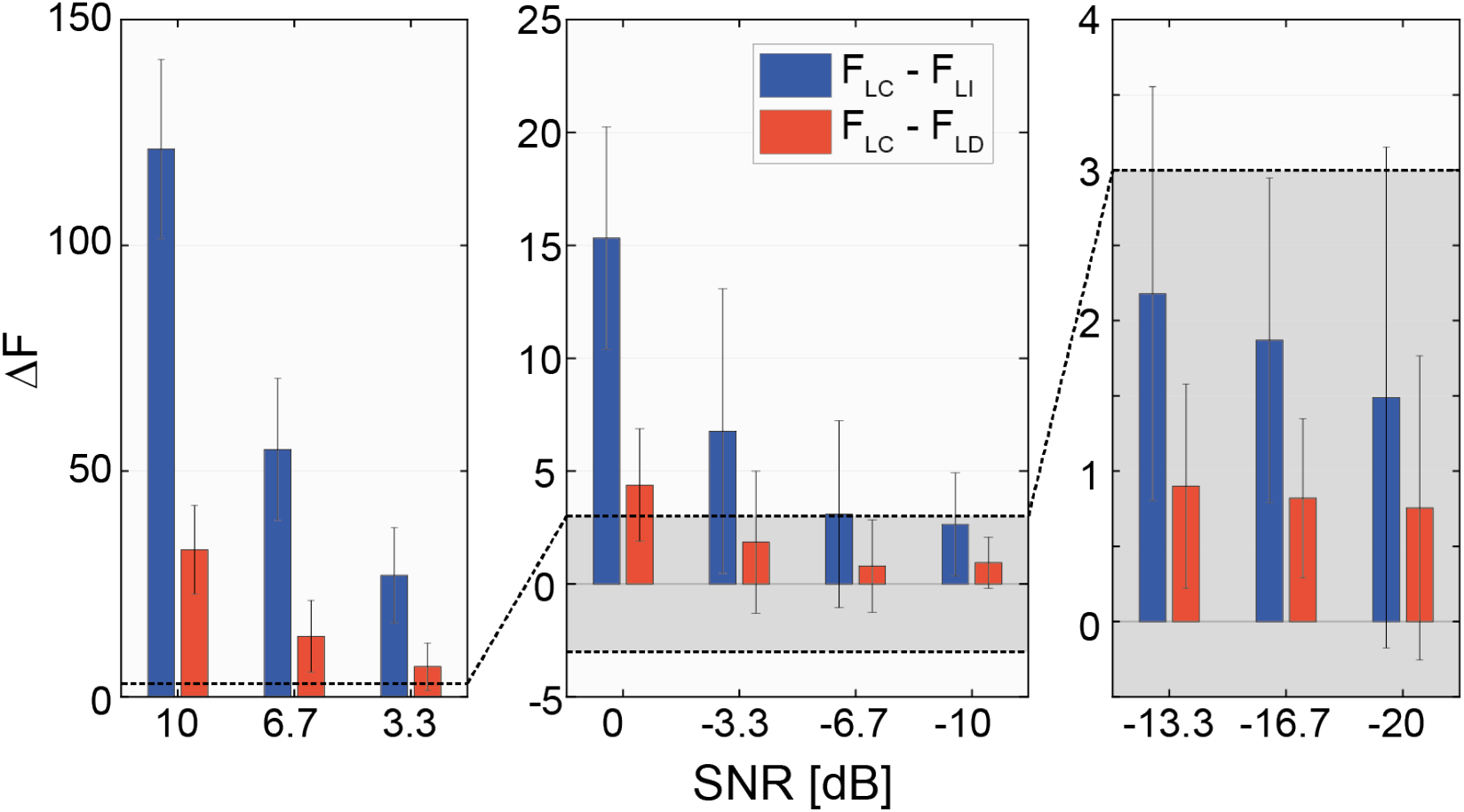
Difference in negative free energy (F) for different SNR levels (x-axis). The results are split into three panels in order to allow for different scaling of the y-axis. The y-axis show differences in F between layer correct (LC, d=2mm) and layer inverted (LI, d=−2mm, blue) and LC and layer detached (LD, d=0mm, red). The LD model does not use any layer specific information. The horizontal dashed line and grey shaded area indicates a difference in F of 3. ΔF>3 is considered strong evidence in favour of the model with the higher F. Note the different scales for the three subfigures. Error bars indicate standard deviations over the 20 simulations.

### Model Comparison: Specificity for Cortical Thickness

Next, we considered the situation where the exact cortical thickness is not known. For this, we created a set of models with thicknesses between −2mm (layer inverted) and 4mm (thick cortex) and tested which of these models best predicted data generated by a model with cortical thickness of 2mm. These simulations were performed for different SNRs of −20, −10, 0, and 10dB, respectively. Figure 5 summarizes the results of these simulations. A discrimination of cortical depth was possible based on the negative free energy difference between models at SNRs of 10dB and 0dB. At a SNR of −10dB and −20dB, none of the models was significantly more likely than the others, i.e. all ΔF < 3. Overall, these simulations illustrate that model comparison can be employed to investigate cortical thickness. However, this requires relatively high SNR of 0dB.

**Figure 5:**
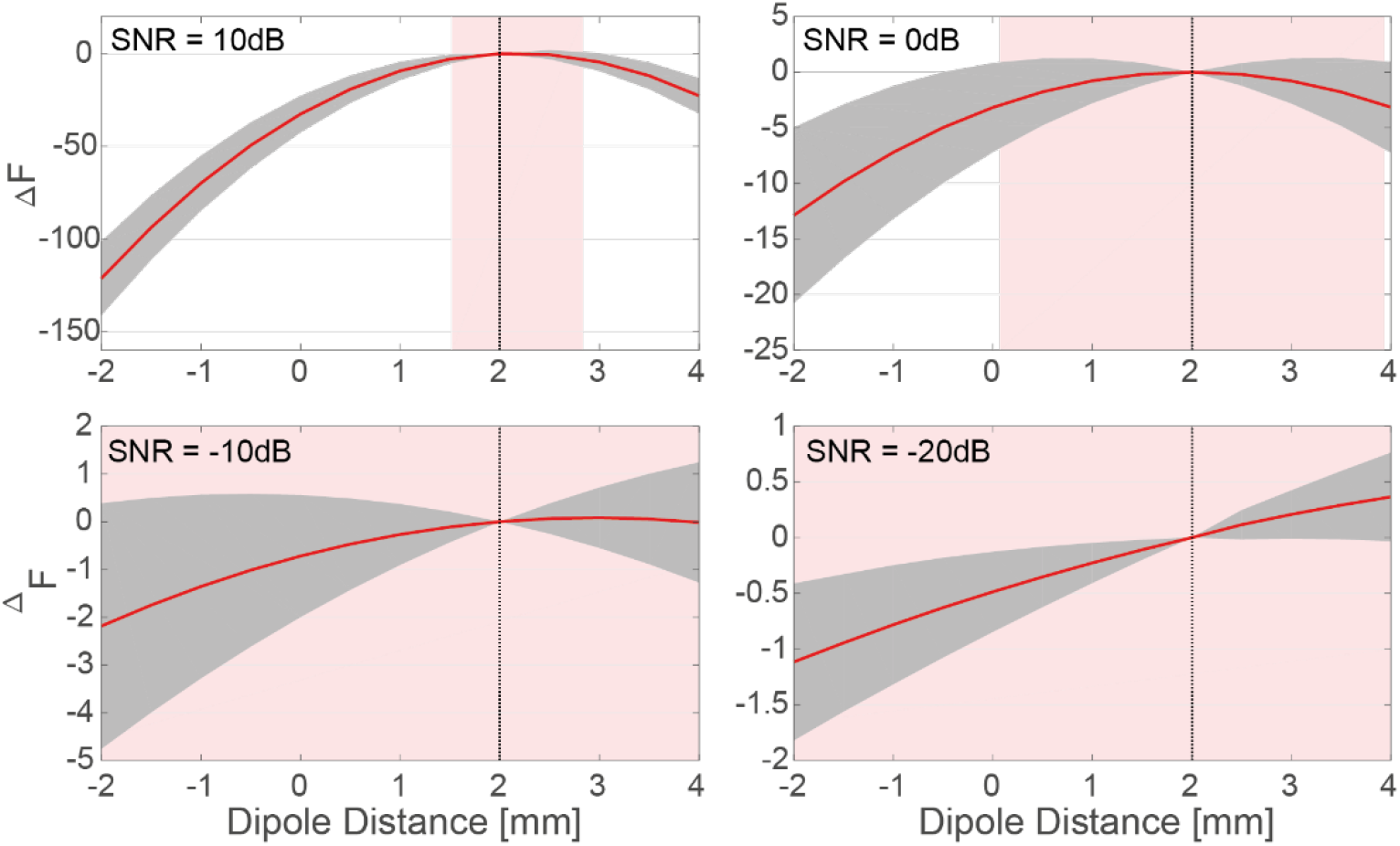
Model comparison of models with different dipole separations. The four subplots show the difference in F when comparing models with differing dipole distance to a model with dipole distance 2 mm that was used to generate the data. SNRs are indicated in the subplots. Red curves show the mean ΔF for 20 simulations of the same SNR. Standard deviations are indicated by the grey shaded area around the curves. Red shaded areas indicate the region where mean ΔF is smaller than 3, i.e. where models cannot be safely distinguished.

**Figure 6:**
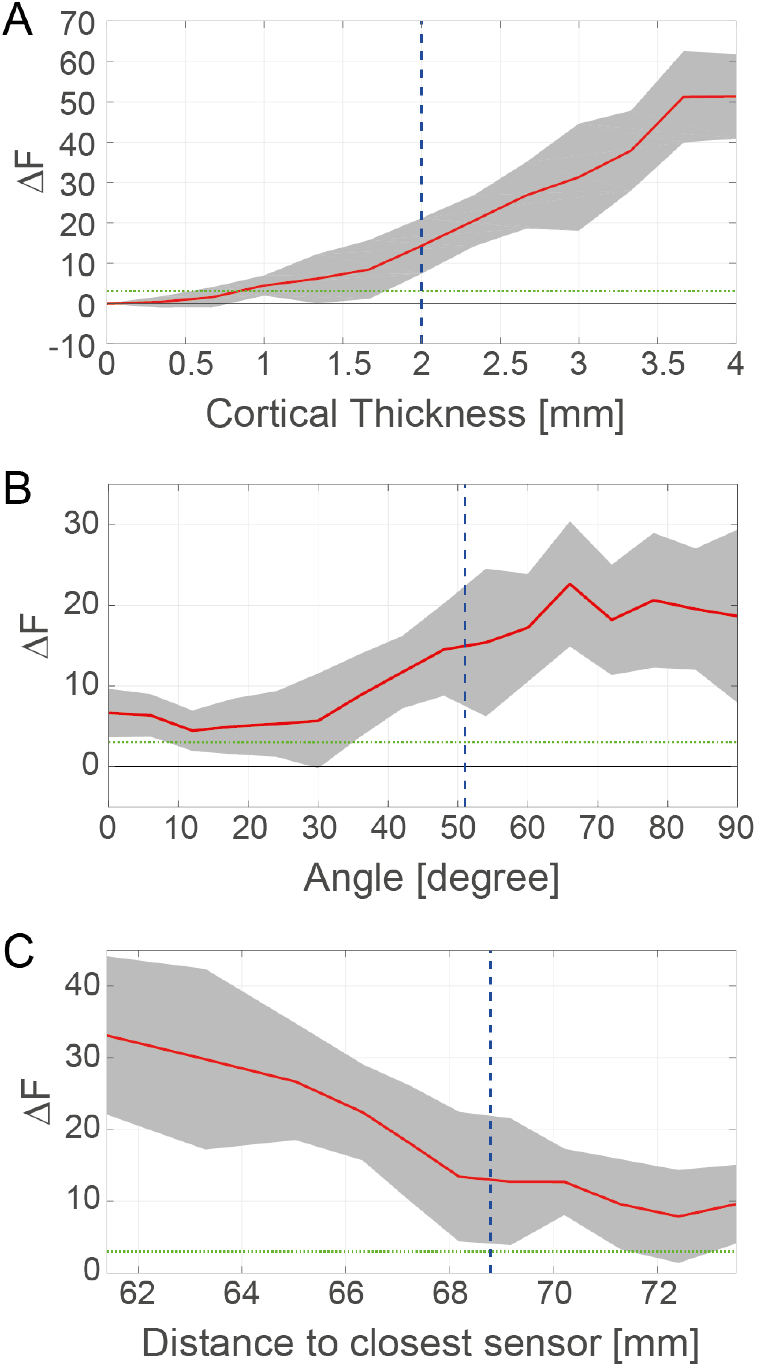
Influence of dipole properties on model selection. The difference in negative free energy (F) between the LC model and the LI model is plotted as a function of **(A)** cortical thickness, i.e. dipole distance, **(B)** orientation of the dipoles, **(C)** distance to the sensor grid. Red traces indicate the mean difference in free energy between the LC and the LI model for data generated by the LC model. Shaded areas are the standard deviations across the 20 simulations performed for each setting. Vertical green dashed lines indicate the values of the main simulations in Figure 4. Horizontal green dotted lines indicate a difference of ΔF = 3.

### Influence of Dipole Properties

In order to determine how properties of the dipole pair impact on a successful layer differentiation, we conducted simulations with certain properties changed. In particular, we varied four different quantities: (i) cortical thickness, (ii) dipole orientation, and (iii) the distance to the sensors, i.e. whether the source was closer to or further away from the skull and, therefore, the closest sensor. For all of these properties, we investigated how much they influenced model discrimination. While in previous simulations the noise was always defined relative to the signal, we adopted a different strategy here. In order to avoid the possibility that changes in signal strength due to changing dipole properties would lead to changes in noise, a fixed noise level was added to all simulations discussed in this paragraph. Concretely, the noise level was chosen to yield an SNR of 0dB when considering the dipole pair used in Figure 4. The same amount of noise was then added to all simulations, potentially leading to different SNRs, e.g. when the current dipole was placed further away from the skull.

Cortical thickness was defined by changing the distance between the deep and superficial dipole for both generation of data as well as model inversion. In simulations, cortical thickness varied from 0 to 4 millimeters. The thicker the cortex, the stronger the LC model was favored over the LI model. For the given noise level, a significant layer discrimination occurred for a cortical thickness above 1 mm. The orientation of the dipole was defined as the angle between the dipole and the radial, i.e. the vector connecting the centre of the volume conductor with the position of the dipole. It was varied from 0° to 90° in steps of 6°. As expected for MEG dipoles the difference in free energy between models increased with increasing angle. In other words, discriminability between layered models is difficult in patches of cortex that are oriented in parallel to the sensor grid (i.e. with current dipoles oriented towards the sensor grid), while sources from sulci where dipoles are oriented in parallel to the sensor grid will facilitate layered model discriminability. While the two first parameters (dipole distance and orientation) correspond mainly to anatomical features of cortex, the distance to the closest sensor can be strongly influenced by how sensors are arranged around the head.

When varying the distance to the closest sensor, the dipole pair location was changed along the line connecting the origin in MNI space with BA3a. The distance between the dipoles (cortical thickness) was set to 2 mm and the dipole angle was kept as in the main simulations. As expected, we observed that the closer the dipole to the sensors, the more successful layer differentiation became. Hence, for discrimination between different layered models, it is highly advantageous to measure close to the sources, i.e. bring the sensors as close as possible to the skull. This would be the case, for example, in MEG measurements using optically pumped magnetometers (OPMs; Tierney et al., 2019).

### Dipole Direction Estimation

In this section, we demonstrate, using simulated data, how the combination of a temporal and spatial model facilitates estimating the correct direction of cortical current flow. This is a long-standing problem in M/EEG as data can often be equally well explained by one of two sources on either side of a sulcus each with different polarity. We assumed that the simulated dipole pair was located in the bank of a sulcus and tested whether it could be distinguished from a second dipole pair at a distance of 7 mm at the opposite side of the sulcus (Fig. 7A). Hence, all dipoles were oriented along the same axis but with different polarities. Data were simulated with the first dipole pair assuming a separation of 2mm and both sources pointing towards the pial surface. We then tested four different models to explain the data. Two of the models were placed at the correct location but with the dipoles pointing either towards the pial surface (pial positive) or away from the pial surface (pial-negative). The other two models were placed at the opposite bank of the sulcus and also had dipoles either oriented towards or away from the pial surface. For all four models, we inverted the data for cortical thicknesses between −4mm (layer inverted) and 4mm.

**Figure 7:**
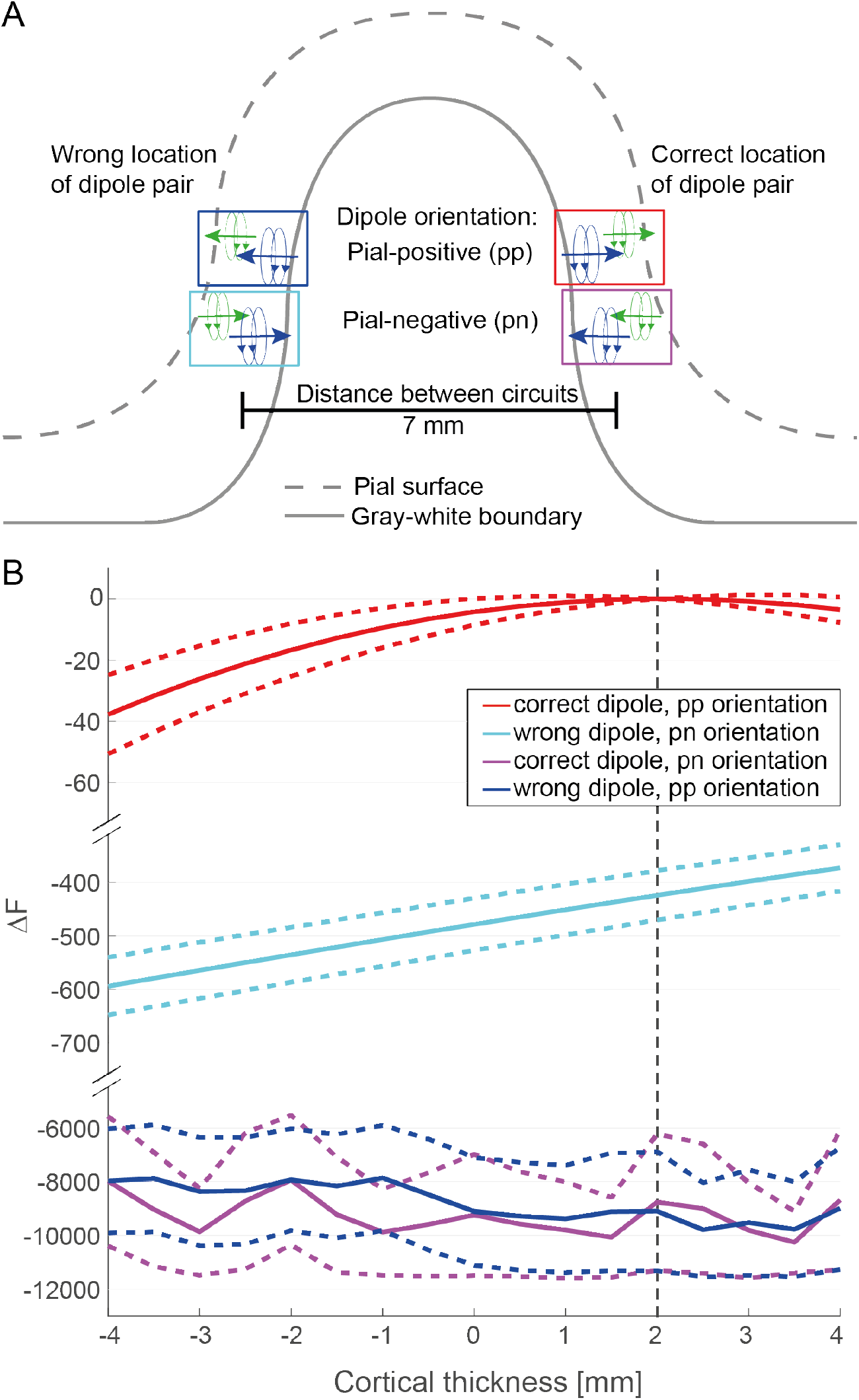
Comparing layered circuits across a sulcus. **A:** Illustration of a simulated scenario. We simulated the problem of distinguishing two layered microcircuits located in the two banks of a sulcus. The correct location used for generating the data was on the right (red and magenta rectangles). The wrong location of the microcircuit was located in the other bank of the sulcus 7 mm away (blue and cyan). For each dipole location we introduce a model with correctly orientated, pial-positive (pp) dipole moments (red and blue) and inverted, pial-negative (pn) dipole moments (magenta and cyan). **B:** Free energy (F) difference of all inverted models compared to the generating model (red, dipole distance: 2mm). Colours as in panel A. Mean (solid lines) free energy difference and standard deviations (dashed lines) over 20 simulations with SNR = 0dB are shown. The dashed vertical line indicates the true (i.e. generating) dipole distance. Note the separation and change in scale between the three segments of the y-axis.

The results of this simulation are depicted in Figure 7. The free energy difference relative to the (true) generating model is shown for all models as a function of the cortical thickness of the estimated model. Models with the correct dipole location and pial-positive dipole orientation performed much better than all other models, independently of the estimated cortical thickness. Examining the competing models more closely, it becomes obvious that a dipole location error of 7mm was much less severe than getting the generator polarity wrong (compare the cyan and violet curves in Figure 7). It is worth noting that for the wrong position, the model with pial-negative dipoles points into the same direction as the generating model (pial-postivie dipoles at true position) because the orientation of cortex is inverted on the other side of the sulcus. As a consequence, the LI model is favoured over the LC model in this scenario. Anatomical information therefore considerably reduces the number of possible source locations as source models situated on half of the sulcal walls will not be fitted well given the temporal model’s prediction. The temporal model can thus act as a constraint to help enforce that the sign of the dipole orientation, in particular of the dominating dipole, has been chosen correctly.

### Reducing the number of sensors

So far, we used data from all 274 sensors for model inversion. While this includes all possible information it makes simulations and model inversion slow. Furthermore, many sensors are quite far away from the dipoles and do not contribute much information because their signal is dominated by noise. For the inversion of real data this could potentially become a problem because a model predicting temporal traces that are 0 everywhere would explain a large part of the data, namely all sensors far away from the current moment. In addition, recent advances in the development of OPMs have led to a new generation of flexible MEG sensors (Boto et al., 2018, 2017) and make it possible to place sensors in specific positions. Here, we investigate how using only a subset of the sensor traces influences model comparison. Simulations were conducted with the standard dipole in BA3 and an SNR of 0 dB as defined above. The results are plotted in Figure 8. We slowly reduced the number N of sensors using two different approaches. First, we always selected the N dipoles with the highest average (between superficial and deep dipole) lead field potential (blue bars in Figure 8). Second, we selected the N dipoles with the largest difference between the two dipole sources (red bars in Figure 8).

**Figure 8:**
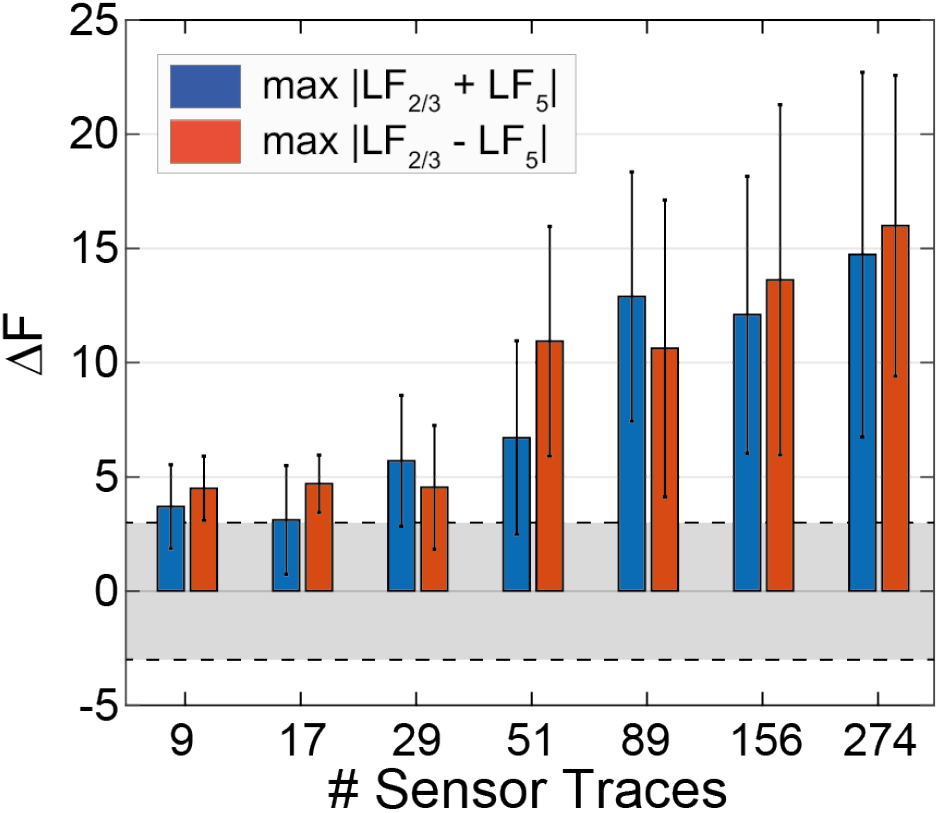
Free energy (F) differences between the LC and LI model using a different number of sensors. The reduced number of N sensors was selected either based on highest summed lead field (blue) or highest difference of lead fields (red) between upper and lower layer. The simulation was conducted at an SNR 0f 0dB.

Generally, free energy differences are larger (and model selection is thus more robust) the more sensors are used. If the sensors with the highest summed lead field are used, average ΔF remains around 3 up to N = 17 sensors. With more than 17 sensors, free energy differences rise clearly above this level. When the sensors with the highest difference between upper and lower layer lead fields are used, free energy differences remain at a level slightly above 3 up to N = 29 and then start to rise. Hence, in order to clearly separate the LC model from the LI model with model selection roughly 30 or more sensors are necessary at an SNR of 0dB. The selection of sensors starts to overlap between the highest average and highest difference when at least 17 sensors are selected. The number of overlapping sensors for different N are: 0 (N=9), 7 (N=17), 15 (N=29), 35 (N=51), 66 (N=89), 120 (N=156) and 274 (N=274). Hence, model discrimination seems to increase when both types of information (average and difference of layer dipoles) are considered.

## Discussion

In this work, we have shown that the contribution of the deep and superficial layers of a cortical column can, in principle, be discriminated using MEG combined with layered electrophysiological models of neural activity. In particular, it is possible to separate the correct layered model that generated the data from a model where the current dipoles of pyramidal neurons are swapped between superficial and deep layers. We further explored how sensitive this model selection is to anatomical details (location and orientation of the dipole) and the distribution of sensors. Overall, our simulations suggest that inference of layered models is feasible in MEG data at a signal to noise ratio that can be achieved with MEG when event related potentials are averaged over several trials. Importantly, we have shown that both temporal and spatial information improve the discrimination between competing models. In particular, the temporal model adds considerable robustness to the spatial model by constraining current flow direction and eliminating competing models on opposing sulcal walls. In the following, we will discuss this approach in the light of alternative methods such as fMRI, highlight its limitations and future improvements, and propose applications on real data.

### Measuring layered cortical activity

Layer-specific activity of evoked potentials can be measured directly with invasive electrodes with many electrical contacts and therefore many electrophysiological measurements throughout cortical depth (Javitt et al., 1996; Schroeder et al., 1995; Self et al., 2013). The resulting local field potentials or current source densities are usually directly compared between different experimental conditions, for example to investigate whether a cortical column is involved in figure ground segregation or not (Self et al., 2013), or to study layered differences of inputs into somatosensory areas (Schroeder et al., 1995). Although these invasive extra-cellular measures are far more “direct” measures than EEG/MEG, they can still be open to interpretation as they are still subject to modelling assumptions (Gratiy et al., 2011; Haegens et al., 2015). Such invasive data were used in conjunction with a two layered DCM (for oscillatory activity) to infer layered activity (Pinotsis et al., 2017). Hence, depth resolving electrophysiological recordings offer an important bridge and testbed for the method proposed here. However, with the exception of patients implanted with electrodes for presurgical localization of epileptic foci, such invasive measures are not possible in humans.

Recent advances in magnetic resonance imaging (MRI) at high-fields (for a review see van der Zwaag et al., 2016) have made it possible to measure functional MRI (fMRI) at submillimeter resolution, allowing for separating at least two cortical layers. While the initial studies focused on early visual (Koopmans et al., 2011, 2010; Siero et al., 2011) and motor areas (Huber et al., 2015; Siero et al., 2011), more recent applications have shown a variety of applications in cognitive neuroscience (De Martino et al., 2015; Finn et al., 2019; Huber et al., 2017; Kok et al., 2016; Lawrence et al., 2019; Muckli et al., 2015). In addition to these cognitive neuroscience approaches, a relationship between oscillatory activity measured with EEG and laminar fMRI has been demonstrated (Scheeringa et al., 2016).

However, there are two major challenges when analyzing and interpreting layered fMRI which measures neural activity only indirectly. First, fMRI activity is filtered by the hemodynamic response function whose dominant frequency is in the range of 0.04 Hz. This might be too slow to fully capture fast computations. Second, layered fMRI can be contaminated by blood draining effects between layers. Although these can be modeled (Havlicek and Uludag, 2019; Heinzle et al., 2016), they pose an additional complexity for the analysis and interpretation of the data.

### Models of layered cortical activity

Computational models of layered cortical activity have a long tradition dating back to the introduction of a canonical microcircuit to model the activity in visual cortex after stimulation of thalamic afferents (Douglas et al., 1989; Douglas and Martin, 1991). This cortical circuit is believed to form the computational substrate for all cortical computations and can, for example, be adapted to model the monkey frontal eye fields that guide eye movements (Heinzle et al., 2007). However, this model was not directly fitted to electrophysiological data. Similarly, the detailed layered compartmental model by Jones and colleagues (Jones et al., 2007) is able to explain evoked potentials as well as oscillatory behaviour (Jones et al., 2009), but was never rigorously fitted to data. A simulation toolbox based on this model allows simulations of EEG and MEG data (Neymotin et al., 2020). In addition, as with most other modelling attempts of EEG/MEG data, the relative laminar displacements of the different neuronal populations were not considered in the forward model. Instead, it was assumed that currents of all pyramidal cells are linearly combined at a virtual point-source.

A similar approach is taken by DCM for EEG/MEG (Kiebel et al., 2009; Moran et al., 2013), where the activity of different cell populations of a canonical microcircuit is summed (with population specific weighting) to yield a dynamically changing electric or current dipole at a single position in cortex. In a recent version of DCM for fMRI (Friston et al., 2019) activity in a layered canonical microcircuit is converted to a single hemodynamic signal. Hence, while both approaches take into account models that consider different layers, the spatial information about layers is not considered in the modelling approach. This is a practical assumption for most EEG/MEG recordings where uncertainty in head or electrode position has a much greater influence on the forward model (Dalal et al., 2014; Hillebrand, 2003). The approach outlined here is a first step towards aligning the advanced temporal modelling literature with more recent spatial modelling work in which relative head-to-sensor geometry issues have been mitigated (Bonaiuto et al., 2018b, 2018a; Troebinger et al., 2014b, 2014b).

### Limitations and potential future improvements

In this section, we discuss limitations of the current approach and suggest some potential improvements for the future. One of the key assumptions of the simulations in this paper was that the location of the dipole pair is known. Future versions could also include a forward model where the geometry of the source configuration is estimated as part of the inversion. This could attempted in a similar manner to conventional DCM, where spatial priors give initial dipole locations (Kiebel et al., 2009, 2006). We should stress that this model contains just a single additional parameter (the displacement) to conventional point source models; and the prior mean and variance of this parameter (related to cortical thickness) can be informed by the anatomy. A second important limitation is that we have so far considered only one region. In most experiments, several cortical regions will be active, and it is not evident how well the mixture of the activity in several cortical microcircuits can be separated into layered activity. While the above points concern mainly the spatial model, one could also think of enhancing the temporal model. One obvious extension would be to use the JCM directly to fit the data. However, this is complicated by the spiking nature of neural activity which is not suitable for the gradient based optimization in the variational Bayes approach used here.

In our simulations, we have used independently and identically distributed Gaussian noise at the sensor level. This is a simplification of the colored and possibly temporally dependent noise in real MEG data (Engemann and Gramfort, 2015). The results indicate that high SNR (−3dB to 0 dB) data is needed to robustly infer layered structure. The estimated SNR of MEG data is roughly in the range of −20 dB to −30dB (Goldenholz et al., 2009) depending on the location of the source. Averaging over 100 or even 1000 trials would bring this range to 0dB to −10dB or 10dB to 0dB, respectively. Indeed, the improved SNR afforded by head-cast MEG allows for identifying laminar-specific spectral responses in sensory and motor cortices (Bonaiuto et al., 2018a). Our approach demonstrates the feasibility for temporal and spatial DCMs for evoked responses, but in the future this approach could be extended to laminar-resolved human MEG of spectral activity.

### Future applications

One of the most prominent contemporary theories of brain function is the Bayesian brain hypothesis (Friston, 2010; Knill and Pouget, 2004). It has been suggested that predictive coding, an implementation of the Bayesian brain, has a natural embedding in cortical microcircuits (Bastos et al., 2012; Mumford, 1992; Rao and Ballard, 1999). Hence, both layered fMRI (Stephan et al., 2019) and layered MEG could provide important experimental evidence in support of or against this hypothesis. One of the advantages of MEG over fMRI is its high temporal resolution. Cortical computation is relatively fast (on the order of tens to hundreds of milliseconds) which is reflected in the timing of event related responses. In order to make inferences about such fast processes, it is highly advantageous to acquire data at a high temporal resolution.

Having established the feasibility of layered DCM in MEG using simulations, the next step will be to invert the model on data from an ERP experiment. Given that the distance to the sensors is critical, the somatosensory cortex seems the most promising candidate. As a next step, one could then move to model several regions. In all these applications, it will be paramount to constrain the spatial location of the dipole pair as narrowly as possible using anatomically precise measurements. One possibility to improve SNR in MEG is the use of OPMs which allow recordings using head-mounted sensors (removing relative head-movement issues), and promise higher SNR recordings as well as potentially much longer recording times (Boto et al., 2018, 2017, 2016). These sensors, placed directly on the scalp, could also take advantage of the higher-spatial frequency information generated by these dipole pairs (figure 2B). We have shown here that layered circuits can be inferred with relatively few cryogenic sensors (roughly 20 in our simulations). This number could potentially be further optimized by using different sensor types of geometries.

## Acknowledgments

This work was supported by the Swiss Data Science Center (SJI), the Biotechnology and Biological Sciences Research Council (BB/M009645/1) (JB), the René and Susanne Braginsky Foundation (KES) and the University of Zurich (KES). The Wellcome Centre for Human Neuroimaging is supported by core funding from Wellcome (203147/Z/16/Z).

## References

Auksztulewicz, R., Blankenburg, F., 2013. Subjective rating of weak tactile stimuli is parametrically encoded in event-related potentials. The Journal of neuroscience: the official journal of the Society for Neuroscience 33, 11878–87. https://doi.org/10.1523/jneurosci.4243-12.2013

Baillet, S., 2017. Magnetoencephalography for brain electrophysiology and imaging. Nature neuroscience 20, 327. https://doi.org/10.1038/nn.4504

Bastos, A.M., Usrey, W.M., Adams, R.A., Mangun, G.R., Fries, P., Friston, K.J., 2012. Canonical microcircuits for predictive coding. Neuron 76, 695–711. https://doi.org/10.1016/j.neuron.2012.10.038

Bonaiuto, J.J., Afdideh, F., Ferez, M., Wagstyl, K., Mattout, J., Bonnefond, M., Barnes, G.R., Bestmann, S., 2019. Estimates of cortical column orientation improve MEG source inversion (preprint). Neuroscience. https://doi.org/10.1101/810267

Bonaiuto, J.J., Meyer, S.S., Little, S., Rossiter, H., Callaghan, M.F., Dick, F., Barnes, G.R., Bestmann, S., 2018a. Lamina-specific cortical dynamics in human visual and sensorimotor cortices. Elife 7. https://doi.org/10.7554/eLife.33977

Bonaiuto, J.J., Rossiter, H.E., Meyer, S.S., Adams, N., Little, S., Callaghan, M.F., Dick, F., Bestmann, S., Barnes, G.R., 2018b. Non-invasive laminar inference with MEG: Comparison of methods and source inversion algorithms. NeuroImage 167, 372–383. https://doi.org/10.1016/j.neuroimage.2017.11.068

Boto, E., Bowtell, R., Kruger, P., Fromhold, T.M., Morris, P.G., Meyer, S.S., Barnes, G.R., Brookes, M.J., 2016. On the Potential of a New Generation of Magnetometers for MEG: A Beamformer Simulation Study. PloS one 11, e0157655. https://doi.org/10.1371/journal.pone.0157655

Boto, E., Holmes, N., Leggett, J., Roberts, G., Shah, V., Meyer, S.S., Munoz, L.D., Mullinger, K.J., Tierney, T.M., Bestmann, S., Barnes, G.R., Bowtell, R., Brookes, M.J., 2018. Moving magnetoencephalography towards real-world applications with a wearable system. Nature 555, 657–661. https://doi.org/10.1038/nature26147

Boto, E., Meyer, S.S., Shah, V., Alem, O., Knappe, S., Kruger, P., Fromhold, T.M., Lim, M., Glover, P.M., Morris, P.G., Bowtell, R., Barnes, G.R., Brookes, M.J., 2017. A new generation of magnetoencephalography: Room temperature measurements using optically-pumped magnetometers. NeuroImage 149, 404–414. https://doi.org/10.1016/j.neuroimage.2017.01.034

Chen, C.C., Henson, R.N., Stephan, K.E., Kilner, J.M., Friston, K.J., 2009. Forward and backward connections in the brain: a DCM study of functional asymmetries. Neuroimage 45, 453–462. https://doi.org/10.1016/j.neuroimage.2008.12.041

Dalal, S.S., Rampp, S., Willomitzer, F., Ettl, S., 2014. Consequences of EEG electrode position error on ultimate beamformer source reconstruction performance. Frontiers in Neuroscience 8. https://doi.org/10.3389/fnins.2014.00042

David, O., Kiebel, S.J., Harrison, L.M., Mattout, J., Kilner, J.M., Friston, K.J., 2006. Dynamic causal modeling of evoked responses in EEG and MEG. NeuroImage 30, 1255–72. https://doi.org/10.1016/j.neuroimage.2005.10.045

De Martino, F., Moerel, M., Ugurbil, K., Goebel, R., Yacoub, E., Formisano, E., 2015. Frequency preference and attention effects across cortical depths in the human primary auditory cortex. Proceedings of the National Academy of Sciences of the United States of America 112, 16036–41. https://doi.org/10.1073/pnas.1507552112

Douglas, R.J., Martin, K.A., 2007. Mapping the matrix: the ways of neocortex. Neuron 56, 226–38. https://doi.org/10.1016/j.neuron.2007.10.017

Douglas, R.J., Martin, K.A., 2004. Neuronal circuits of the neocortex. Annual review of neuroscience 27, 419–51. https://doi.org/10.1146/annurev.neuro.27.070203.144152

Douglas, R.J., Martin, K.A., 1991. A functional microcircuit for cat visual cortex. J Physiol (Lond) 440, 735–769.

Douglas, R.J., Martin, K.A.C., Whitteridge, D., 1989. A Canonical Microcircuit for Neocortex. Neural Computation 1, 480–488.

Engemann, D.A., Gramfort, A., 2015. Automated model selection in covariance estimation and spatial whitening of MEG and EEG signals. Neuroimage 108, 328–342. https://doi.org/10.1016/j.neuroimage.2014.12.040

Finn, E.S., Huber, L., Jangraw, D.C., Molfese, P.J., Bandettini, P.A., 2019. Layer-dependent activity in human prefrontal cortex during working memory. Nat. Neurosci. 22, 1687–1695. https://doi.org/10.1038/s41593-019-0487-z

Fischl, B., Dale, A.M., 2000. Measuring the thickness of the human cerebral cortex from magnetic resonance images. Proceedings of the National Academy of Sciences of the United States of America 97, 11050–5. https://doi.org/10.1073/pnas.200033797

Friston, K., 2010. The free-energy principle: a unified brain theory? Nature reviews. Neuroscience 11, 127–38. https://doi.org/10.1038/nrn2787

Friston, K., 2005. A theory of cortical responses. Philosophical transactions of the Royal Society of London. Series B, Biological sciences 360, 815–36. https://doi.org/10.1098/rstb.2005.1622

Friston, K., Mattout, J., Trujillo-Barreto, N., Ashburner, J., Penny, W., 2007. Variational free energy and the Laplace approximation. NeuroImage 34, 220–34. https://doi.org/10.1016/j.neuroimage.2006.08.035

Friston, K.J., Preller, K.H., Mathys, C., Cagnan, H., Heinzle, J., Razi, A., Zeidman, P., 2019. Dynamic causal modelling revisited. NeuroImage 199, 730–744. https://doi.org/10.1016/j.neuroimage.2017.02.045

Garrido, M.I., Friston, K.J., Kiebel, S.J., Stephan, K.E., Baldeweg, T., Kilner, J.M., 2008. The functional anatomy of the MMN: a DCM study of the roving paradigm. NeuroImage 42, 936–44. https://doi.org/10.1016/j.neuroimage.2008.05.018

Goldenholz, D.M., Ahlfors, S.P., Hamalainen, M.S., Sharon, D., Ishitobi, M., Vaina, L.M., Stufflebeam, S.M., 2009. Mapping the signal-to-noise-ratios of cortical sources in magnetoencephalography and electroencephalography. Human brain mapping 30, 1077–86. https://doi.org/10.1002/hbm.20571

Gratiy, S.L., Devor, A., Einevoll, G.T., Dale, A.M., 2011. On the estimation of population-specific synaptic currents from laminar multielectrode recordings. Front Neuroinform 5, 32. https://doi.org/10.3389/fninf.2011.00032

Haegens, S., Barczak, A., Musacchia, G., Lipton, M.L., Mehta, A.D., Lakatos, P., Schroeder, C.E., 2015. Laminar Profile and Physiology of the α Rhythm in Primary Visual, Auditory, and Somatosensory Regions of Neocortex. J. Neurosci. 35, 14341–14352. https://doi.org/10.1523/JNEUROSCI.0600-15.2015

Havlicek, M., Uludag, K., 2019. A dynamical model of the laminar BOLD response (preprint). Neuroscience. https://doi.org/10.1101/609099

Heinzle, J., Hepp, K., Martin, K.A., 2007. A microcircuit model of the frontal eye fields. The Journal of neuroscience: the official journal of the Society for Neuroscience 27, 9341–53. https://doi.org/10.1523/JNEUROSCI.0974-07.2007

Heinzle, J., Koopmans, P.J., den Ouden, H.E., Raman, S., Stephan, K.E., 2016. A hemodynamic model for layered BOLD signals. NeuroImage 125, 556–70. https://doi.org/10.1016/j.neuroimage.2015.10.025

Hillebrand, A., 2003. The use of anatomical constraints with MEG beamformers. NeuroImage 20, 2302–2313. https://doi.org/10.1016/j.neuroimage.2003.07.031

Huber, L., Goense, J., Kennerley, A.J., Trampel, R., Guidi, M., Reimer, E., Ivanov, D., Neef, N., Gauthier, C.J., Turner, R., Moller, H.E., 2015. Cortical lamina-dependent blood volume changes in human brain at 7T. NeuroImage 107, 23–33. https://doi.org/10.1016/j.neuroimage.2014.11.046

Huber, L., Handwerker, D.A., Jangraw, D.C., Chen, G., Hall, A., Stüber, C., Gonzalez-Castillo, J., Ivanov, D., Marrett, S., Guidi, M., Goense, J., Poser, B.A., Bandettini, P.A., 2017. High-Resolution CBV-fMRI Allows Mapping of Laminar Activity and Connectivity of Cortical Input and Output in Human M1. Neuron 96, 1253–1263.e7. https://doi.org/10.1016/j.neuron.2017.11.005

Jansen, B.H., Rit, V.G., 1995. Electroencephalogram and visual evoked potential generation in a mathematical model of coupled cortical columns. Biological cybernetics 73, 357–66.

Javitt, D.C., Steinschneider, M., Schroeder, C.E., Arezzo, J.C., 1996. Role of cortical N-methyl-D-aspartate receptors in auditory sensory memory and mismatch negativity generation: implications for schizophrenia. Proceedings of the National Academy of Sciences of the United States of America 93, 11962–7.

Jones, S.R., Pritchett, D.L., Sikora, M.A., Stufflebeam, S.M., Hämäläinen, M., Moore, C.I., 2009. Quantitative Analysis and Biophysically Realistic Neural Modeling of the MEG Mu Rhythm: Rhythmogenesis and Modulation of Sensory-Evoked Responses. Journal of neurophysiology 102, 3554–3572. https://doi.org/10.1152/jn.00535.2009

Jones, S.R., Pritchett, D.L., Stufflebeam, S.M., Hamalainen, M., Moore, C.I., 2007. Neural correlates of tactile detection: a combined magnetoencephalography and biophysically based computational modeling study. The Journal of neuroscience: the official journal of the Society for Neuroscience 27, 10751–64. https://doi.org/10.1523/JNEUROSCI.0482-07.2007

Kass, R.E., Raftery, A.E., 1995. Bayes Factors. J Am Stat Assoc 90, 773–795. https://doi.org/Doi10.2307/2291091

Kiebel, S.J., David, O., Friston, K.J., 2006. Dynamic causal modelling of evoked responses in EEG/MEG with lead field parameterization. NeuroImage 30, 1273–84. https://doi.org/10.1016/j.neuroimage.2005.12.055

Kiebel, S.J., Garrido, M.I., Moran, R., Chen, C.C., Friston, K.J., 2009. Dynamic causal modeling for EEG and MEG. Human brain mapping 30, 1866–76. https://doi.org/10.1002/hbm.20775

Knill, D.C., Pouget, A., 2004. The Bayesian brain: the role of uncertainty in neural coding and computation. Trends in Neurosciences 27, 712–9. https://doi.org/10.1016/j.tins.2004.10.007

Kok, P., Bains, L.J., van Mourik, T., Norris, D.G., de Lange, F.P., 2016. Selective Activation of the Deep Layers of the Human Primary Visual Cortex by Top-Down Feedback. Current biology: CB 26, 371–6. https://doi.org/10.1016/j.cub.2015.12.038

Koopmans, P.J., Barth, M., Norris, D.G., 2010. Layer-specific BOLD activation in human V1. Human brain mapping 31, 1297–304. https://doi.org/10.1002/hbm.20936

Koopmans, P.J., Barth, M., Orzada, S., Norris, D.G., 2011. Multi-echo fMRI of the cortical laminae in humans at 7 T. NeuroImage 56, 1276–85. https://doi.org/10.1016/j.neuroimage.2011.02.042

Lawrence, S.J., Norris, D.G., de Lange, F.P., 2019. Dissociable laminar profiles of concurrent bottom-up and top-down modulation in the human visual cortex. eLife 8, e44422. https://doi.org/10.7554/eLife.44422

Litvak, V., Mattout, J., Kiebel, S., Phillips, C., Henson, R., Kilner, J., Barnes, G., Oostenveld, R., Daunizeau, J., Flandin, G., Penny, W., Friston, K., 2011. EEG and MEG data analysis in SPM8. Computational intelligence and neuroscience 2011, 852961. https://doi.org/10.1155/2011/852961

Moran, R., Pinotsis, D.A., Friston, K., 2013. Neural masses and fields in dynamic causal modeling. Frontiers in computational neuroscience 7, 57. https://doi.org/10.3389/fncom.2013.00057

Muckli, L., De Martino, F., Vizioli, L., Petro, L.S., Smith, F.W., Ugurbil, K., Goebel, R., Yacoub, E., 2015. Contextual Feedback to Superficial Layers of V1. Current biology: CB 25, 2690–5. https://doi.org/10.1016/j.cub.2015.08.057

Mumford, D., 1992. On the computational architecture of the neocortex. II. The role of cortico-cortical loops. Biological cybernetics 66, 241–51.

Neymotin, S.A., Daniels, D.S., Caldwell, B., McDougal, R.A., Carnevale, N.T., Jas, M., Moore, C.I., Hines, M.L., Hämäläinen, M., Jones, S.R., 2020. Human Neocortical Neurosolver (HNN), a new software tool for interpreting the cellular and network origin of human MEG/EEG data. Elife 9. https://doi.org/10.7554/eLife.51214

Nolte, G., 2003. The magnetic lead field theorem in the quasi-static approximation and its use for magnetoencephalography forward calculation in realistic volume conductors. Physics in medicine and biology 48, 3637–52.

Papadelis, C., Eickhoff, S.B., Zilles, K., Ioannides, A.A., 2011. BA3b and BA1 activate in a serial fashion after median nerve stimulation: Direct evidence from combining source analysis of evoked fields and cytoarchitectonic probabilistic maps. NeuroImage 54, 60–73. https://doi.org/10.1016/j.neuroimage.2010.07.054

Pinotsis, D.A., Geerts, J.P., Pinto, L., FitzGerald, T.H., Litvak, V., Auksztulewicz, R., Friston, K.J., 2017. Linking canonical microcircuits and neuronal activity: Dynamic causal modelling of laminar recordings. NeuroImage 146, 355–366. https://doi.org/10.1016/j.neuroimage.2016.11.041

Rao, R.P., Ballard, D.H., 1999. Predictive coding in the visual cortex: a functional interpretation of some extra-classical receptive-field effects. Nature neuroscience 2, 79–87. https://doi.org/10.1038/4580

Scheeringa, R., Koopmans, P.J., van Mourik, T., Jensen, O., Norris, D.G., 2016. The relationship between oscillatory EEG activity and the laminar-specific BOLD signal. Proc Natl Acad Sci USA 113, 6761–6766. https://doi.org/10.1073/pnas.1522577113

Scholtens, L.H., de Reus, M.A., van den Heuvel, M.P., 2015. Linking contemporary high resolution magnetic resonance imaging to the von Economo legacy: A study on the comparison of MRI cortical thickness and histological measurements of cortical structure. Human brain mapping 36, 3038–46. https://doi.org/10.1002/hbm.22826

Schroeder, C.E., Seto, S., Arezzo, J.C., Garraghty, P.E., 1995. Electrophysiological evidence for overlapping dominant and latent inputs to somatosensory cortex in squirrel monkeys. Journal of neurophysiology 74, 722–32. https://doi.org/10.1152/jn.1995.74.2.722

Self, M.W., van Kerkoerle, T., Super, H., Roelfsema, P.R., 2013. Distinct roles of the cortical layers of area V1 in figure-ground segregation. Current biology: CB 23, 2121–9. https://doi.org/10.1016/j.cub.2013.09.013

Shipp, S., 2016. Neural Elements for Predictive Coding. Frontiers in psychology 7, 1792. https://doi.org/10.3389/fpsyg.2016.01792

Siero, J.C., Petridou, N., Hoogduin, H., Luijten, P.R., Ramsey, N.F., 2011. Cortical depth-dependent temporal dynamics of the BOLD response in the human brain. Journal of cerebral blood flow and metabolism: official journal of the International Society of Cerebral Blood Flow and Metabolism 31, 1999–2008. https://doi.org/10.1038/jcbfm.2011.57

Stephan, K.E., Petzschner, F.H., Kasper, L., Bayer, J., Wellstein, K.V., Stefanics, G., Pruessmann, K.P., Heinzle, J., 2019. Laminar fMRI and computational theories of brain function. NeuroImage 197, 699–706. https://doi.org/10.1016/j.neuroimage.2017.11.001

Tierney, T.M., Holmes, N., Mellor, S., López, J.D., Roberts, G., Hill, R.M., Boto, E., Leggett, J., Shah, V., Brookes, M.J., Bowtell, R., Barnes, G.R., 2019. Optically pumped magnetometers: From quantum origins to multi-channel magnetoencephalography. Neuroimage 199, 598–608. https://doi.org/10.1016/j.neuroimage.2019.05.063

Troebinger, L., Lopez, J.D., Lutti, A., Bestmann, S., Barnes, G., 2014a. Discrimination of cortical laminae using MEG. NeuroImage 102 Pt 2, 885–93. https://doi.org/10.1016/j.neuroimage.2014.07.015

Troebinger, L., Lopez, J.D., Lutti, A., Bradbury, D., Bestmann, S., Barnes, G., 2014b. High precision anatomy for MEG. NeuroImage 86, 583–91. https://doi.org/10.1016/j.neuroimage.2013.07.065

van der Zwaag, W., Schafer, A., Marques, J.P., Turner, R., Trampel, R., 2016. Recent applications of UHF-MRI in the study of human brain function and structure: a review. NMR in biomedicine 29, 1274–88. https://doi.org/10.1002/nbm.3275

